# Simultaneous observation of RNAs and their binding proteins in plant cells

**DOI:** 10.1101/2024.09.27.615539

**Authors:** Junfei Ma, Yunhan Wang, Svetlana Y. Folimonova, Bin Liu, Ying Wang

**Author notes:** Correspondence: Junfei Ma and Ying Wang, (JM) and (YW). These authors contributed equally to the work (Junfei Ma, Yunhan Wang).

## Abstract

RNA is key component of a versatile group of complexes that play diverse roles in a wide range of biological processes, including plant acclimation to ever-changing environments. From synthesis to degradation, RNA molecules interact with cognate proteins that assist in processes such as transcription, splicing, modification, trafficking, and the execution of their functions. While numerous valuable techniques exist to study RNA-protein interactions, visualizing RNAs and their associated proteins simultaneously within cells remains challenging, despite its potential to provide deeper insights into the biology of RNA-protein interactions. In this study, we adapted a modified immunofluorescence (IF) assay in combination with RNA fluorescence in situ hybridization (FISH) to successfully visualize the colocalization of endogenous as well as infectious RNAs with their cognate binding proteins in plant nucleus. This new method that combines IF and FISH will facilitate future studies on RNA and protein colocalization in various plant systems.

## Introduction

Mounting evidence has highlighted the essential roles of diverse classes of RNAs across a wide range of biological processes (Sharp, 2009). From transcription to decay, RNAs interact with specific RNA-binding proteins, and these interactions often correlate with distinct biological functions (Street et al., 2024). In certain cases, RNA-protein complexes organize into membrane-less organelles via liquid-liquid phase separation (Miao, Chodasiewicz, & Fang, 2024). Examples include Processing bodies and Stress Granules involved in mRNA turnover or storage (Chantarachot & Bailey-Serres, 2018), SERRATE-driven dicing bodies in microRNA processing (Xie et al., 2021), FLL2-promoted phase separation of polyadenylation complexes for mRNA 3’ end processing (Fang et al., 2019), etc. Although tools such as RNA immunoprecipitation and gel electrophoresis mobility shift assays are valuable for studying RNA-protein interactions, observing RNA subcellular localization or simultaneously visualizing specific RNAs with their associated proteins in cells remains challenging, hindering the understanding of RNA-protein dynamics in the context of cell biology.

The MS2-MCP system enables live-cell imaging of RNAs by using the MS2-binding coat protein (MCP) tagged with a green fluorescent protein (GFP) to recognize RNAs fused with MS2 hairpins (Syed & Lim, 2024). By tracking MSP-fused GFP fluorescence, RNA subcellular localization can be observed in living cells with high specificity and temporal resolution. RNA-binding proteins can be co-visualized by tagging them with fluorescent proteins other than GFP. In addition to the MS2-MCP system, RNA fluorescence aptamers also allow visualization of fusion RNAs in living cells (Sachdeva, Garg, Godding, Way, & Silver, 2014). However, both systems pose a risk of disrupting RNA overall structures and functions. For example, any insertion in the viroid RNA genome can lead to a loss of infectivity, limiting the applications of these methods to certain RNAs.

Observing proteins and RNAs in chemically fixed cells can provide insights into their subcellular dynamics as well. Immunofluorescence (IF) detects specific proteins, while RNA fluorescence in situ hybridization (FISH) reveals RNA subcellular localization patterns. The development of single-molecule RNA FISH (smFISH) enables the quantitative observation of mRNAs, but it requires dozens of short, specific oligos (17-22 nt) and may not be suitable for short RNAs or those with limited unique sequences (Duncan, Johansson, & Ding, 2023). Moreover, traditional IF and FISH are often incompatible, because the formamide used in FISH tend to mask antigens, hindering protein detection (B. Ma & Tanese, 2013; Meng, Zhao, & Lao, 2018; Meyer, Garzia, & Tuschl, 2017). Notably, some reports showed that fluorescent proteins (FPs) could retain fluorescence activity to outline protein subcellular localization after RNA FISH steps (Duncan et al., 2023; Huang, Guillotin, Rahni, Birnbaum, & Wagner, 2023; Zhao, Fonseca, Meschichi, Sicard, & Rosa, 2023). However, this approach is mostly limited to observing proteins colocalized with cytosolic RNAs because RNA-FISH protocols for observing nuclear RNAs often denature FPs by high temperatures and stringent chemical treatments (Alkaabi, Yafea, & Ashraf, 2005; Jolly, Mongelard, Robert-Nicoud, & Vourc’h, 1997; Qi & Ding, 2003; Yokoyama, Ogawa, Nozu, & Hashimoto, 1990).

To address this shortcoming, we developed a new strategy that combines IF and FISH to simultaneously visualize an RNA and its interacting protein partner in same cells. This approach is straightforward, does not require extensive cloning or transgenic plants (if a specific primary antibody is available), and reliably detects RNAs and proteins in the nucleus. We successfully tested the method by using a classic RNA-binding protein fibrillarin and one of its RNA partners (U3 small nucleolar RNA/snoRNA), which often function in biomolecular condensates for processing nascent rRNAs (Yao et al., 2019).

We also applied this method to examining viroid-host interactions at the cellular and subcellular levels. Viroids are single-stranded, circular, noncoding RNA pathogens that primarily infect crops (J. Ma, Dissanayaka Mudiyanselage, Hao, & Wang, 2023; Ortola & Daros, 2023). There are two viroid families, *Pospiviroidae* and *Avsunviroidae*. Members of *Pospiviroidae* replicate in the nucleus using host RNA polymerase II (Pol II)(Wang, 2021). Unlike the well-known 12-subunit Pol II, a modified Pol II with 6-7 subunits transcribes the potato spindle tuber viroid (PSTVd) RNA (Dissanayaka Mudiyanselage et al., 2022), assisted by an RNA-specific transcription factor (TFIIIA-7ZF)(Dissanayaka Mudiyanselage et al., 2022; Wang et al., 2016). Despite the progress achieved in the understanding of the PSTVd replication, the detailed subcellular localization and the pattern of Pol II transcription on PSTVd RNA remain unclear. Using our new method, we observed the nuclear distribution patterns of Pol II (represented by RPB1) and PSTVd sense-strand RNA in protoplasts. Therefore, this method for simultaneous observation of RNAs and proteins in plant cells is valuable for studying various nuclear-localized noncoding RNAs and their associated proteins, including those involved in liquid-liquid phase separation across diverse biological processes.

## Results

### Optimizing RNA FISH protocol for detecting RNA in the nucleus

First, we sought to use PSTVd RNA as an example to optimize the procedures for detecting RNA in fixed plant cells. Notably, a detailed protocol for this type of experiments is not currently available. Due to PSTVd’s relatively short length (359 nt), designing dozens of oligo probes, as required in some single-molecule RNA FISH protocols, is not possible. Instead, we used traditional *in vitro* transcription method to generate RNA oligos.

The key factor is optimizing probe size. We designed two probes around 100 nt in length that are short enough to facilitate nuclear entry. Each probe contained a maximum of 25-30 AlexaFluor488-labeled UTPs. We used both probes, achieving up to 54 labeled UTPs per PSTVd RNA to enhance the signal-to-background ratio. We tested two general protocols reported in previous RNA FISH experiments that observed nuclear PSTVd accumulation (Qi & Ding, 2002; Yokoyama et al., 1990). Interestingly, combining the penetration steps from both protocols yielded better, more consistent results (Figure 1). PSTVd displayed a unique distribution pattern, with a dominant signal near the nucleolar region, as indicated by the hollow DAPI staining (Figure 1). This PSTVd RNA condensate pattern is consistent with previous reports (Qi & Ding, 2002, 2003). Notably, this protocol also worked well for the endogenous U3 snoRNA as we performed colocalization analysis of U3 snoRNA and fibrillarin (Figure 2A).

**Figure 1.**
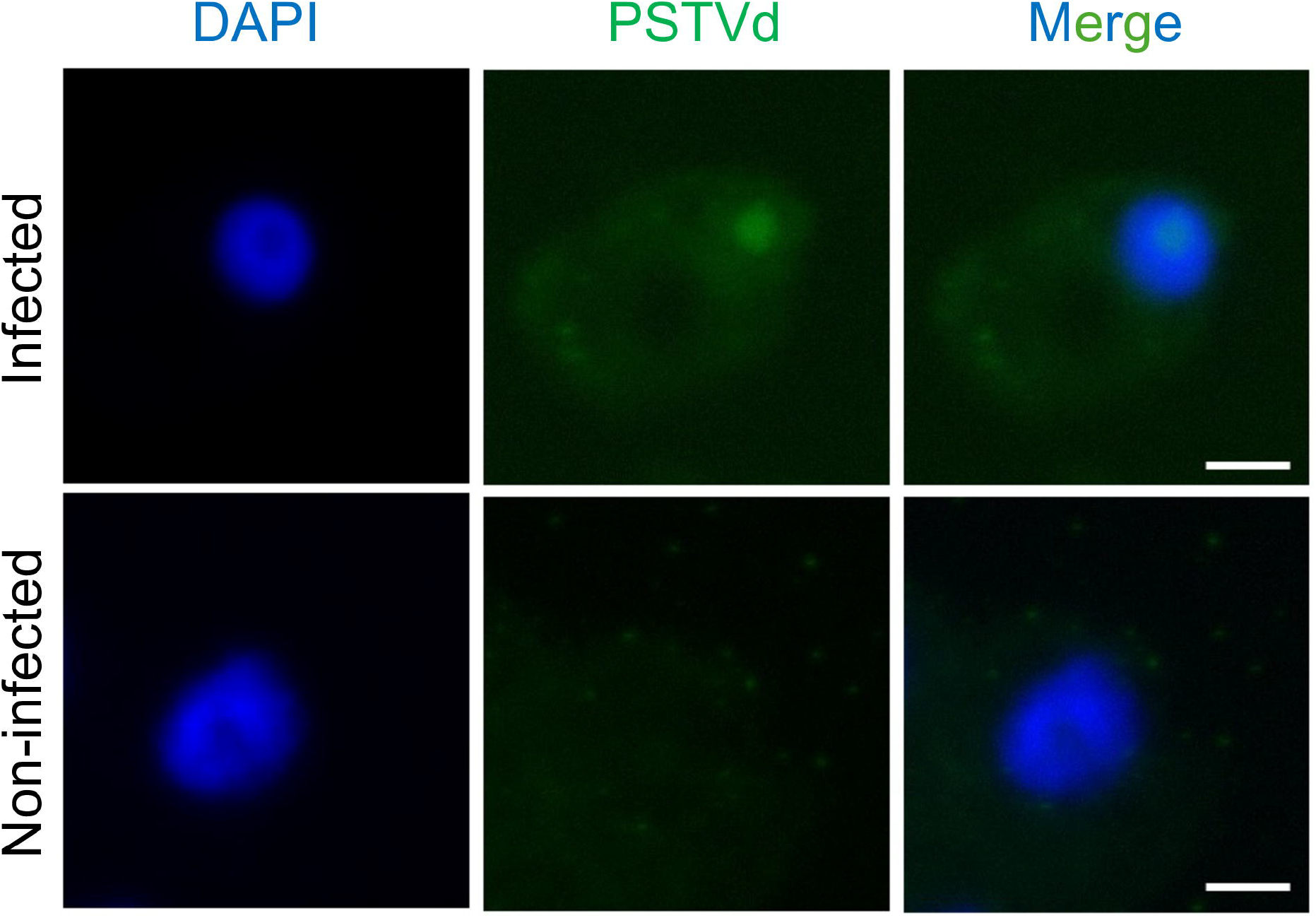
An optimized RNA FISH protocol for observing RNA in plant nucleus. Nucleolar-enriched potato spindle tuber viroid (PSTVd) detected by Alexa Fluor488-labeled riboprobe. Scale bar, 8 μm.

**Figure 2.**
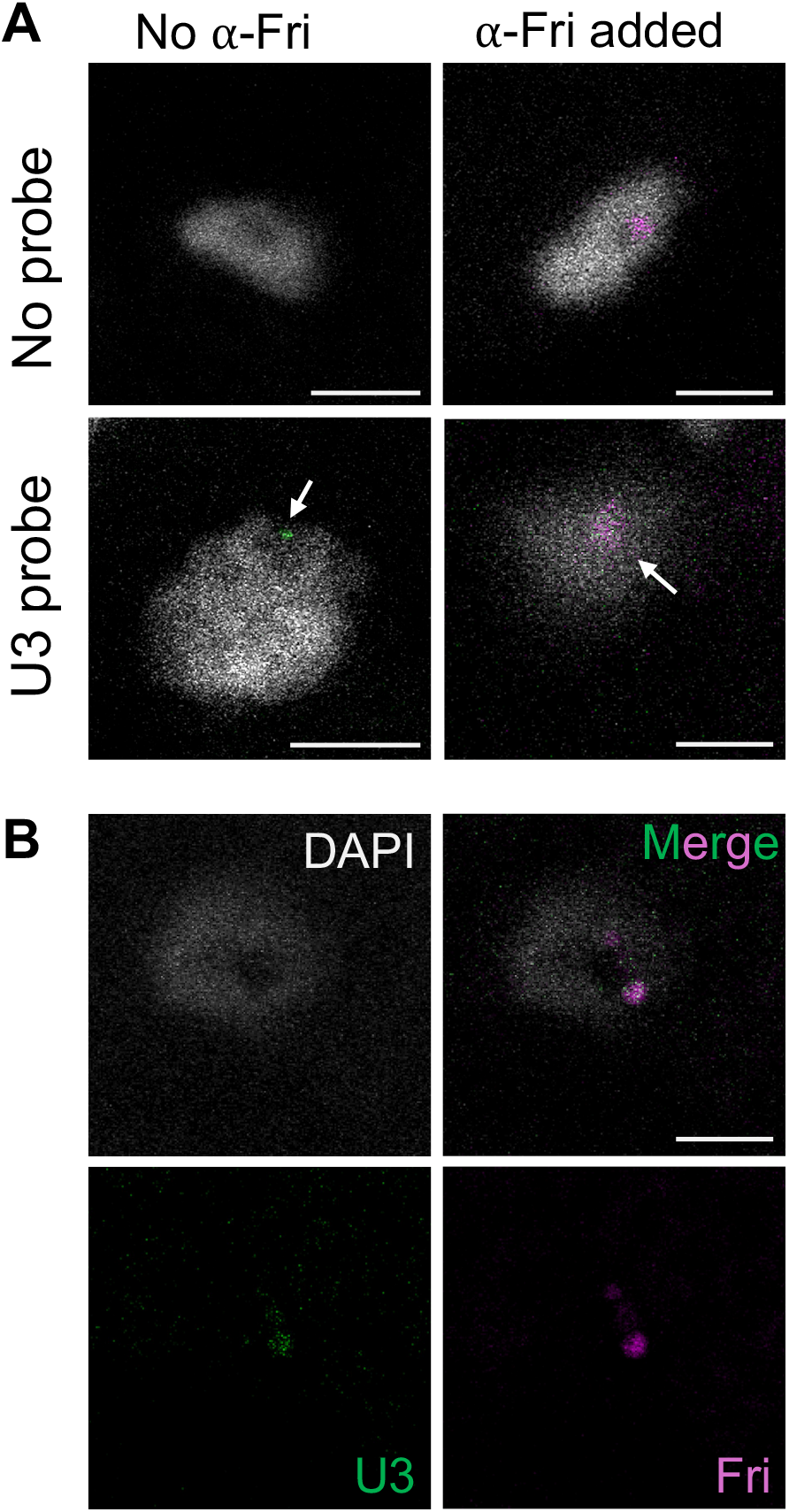
Colocalization of fibrillarin and U3 snoRNA. A) fibrillarin (Fri) is detected by ⍰-Fri1 monoclonal antibody and Alexa Fluor594-labeled anti-mouse secondary antibody (magenta color). Secondary antibody was applied to all treatments. U3 snoRNA is detected by Alexa Fluor488-labeled specific riboprobe (green). White arrow heads point to some colocalized loci. B) Separate color channels of a representative sample showing the colocalization of U3 snoRNA and fibrillarin. Scale bar, 5 μm.

### Modifying the traditional immunofluorescence IF protocol allowing simultaneous observation of fibrillarin and U3 small nucleolar RNA

Only a limited number of immunofluorescence protocols are available for plant samples. Based on a previous protocol (Borg, Buendia, & Berger, 2019), we performed extensive trials by using fibrillarin as a target and established one that works well for this endogenous protein. Fibrillarin is an RNA-binding protein that associates with many endogenous as well as infectious RNAs, such as U3, U8, and U13 small nucleolar RNAs (snoRNAs) (Tyc & Steitz, 1989), ribosomal RNAs (rRNAs) (Rakitina, Taliansky, Brown, & Kalinina, 2011), ELF18-INDUCED LONG NONCODING RNA 1 (Seo, Diloknawarit, Park, & Chua, 2019), tobacco mosaic virus (Rakitina et al., 2011), bamboo mosaic virus satellite RNA (Chang et al., 2016), etc. Fibrillarin is a methyltransferase enzyme that is critical for rRNAs’ processing (Rodriguez-Corona, Sobol, Rodriguez-Zapata, Hozak, & Castano, 2015), and U3 snoRNA-containing ribonucleoprotein guides fibrillarin to perform the first step of pre-ribosomal RNA processing (Kass, Tyc, Steitz, & Sollner-Webb, 1990). To allow simultaneous observation of U3 snoRNA and fibrillarin in the nucleolus, we slightly modified the IF protocol to crosslink the samples twice, pre-IF crosslinking and post-IF crosslinking. The purpose is to preserve the antibody-antigen interaction in harsh RNA-FISH procedures. As shown in Figure 2A, we successfully observed a distinct signal of fibrillarin in the nucleolus.

Based on our RNA FISH protocol, we designed a full-length U3 snoRNA-specific probe, which allow maximum of 38 AlexaFluor488-labeled UTPs. As shown in Figure 2A, U3 snoRNA specific signals were observed in the nucleolus and often co-localized with fibrillarin when both specific antibodies and probes were applied. Separate color channels of another example also confirmed the colocalization around the nucleolar region (Figure 2B). This colocalization pattern resembles previous observations in HeLa cells (Leary, Terns, & Huang, 2004). We analyzed the RNA-protein colocalization by using the JACoP plugin of Image J (Bolte & Cordelieres, 2006) and found the Pearson’s co-efficiency values range from 0.735 to 0.949 (Figure S1), an indication of co-localization.

### An alternative immunofluorescence (IF) approach allowing simultaneous observation of PSTVd and Pol II

With the success of observing the co-localization of U3 snoRNA and fibrillarin, we aimed to observe the subcellular localization of Pol II and PSTVd, which would deepen our understanding of plant-viroid interactions. However, our initial attempts using the above-mentioned IF protocol were unsuccessful mainly due to the failure of antibody penetration into the fixed nuclei. Inspired by a recent study on RNase L-induced bodies that directly incubate antibodies with culture cells (Watkins & Burke, 2024), we reasoned that incubating antibodies directly with protoplasts could be an effective alternative IF approach. We tested this approach by observing GFP protein localization in the *Nicotiana benthamiana* 16C line, which expresses a GFP5 variant with an endoplasmic reticulum localization signal (Philips et al., 2017). Figure 3 shows the GFP fluorescence signal in the live protoplasts as well as in the cells incubated with the antibodies. Signals from the Alexa Fluor594-conjugated anti-GFP antibody largely highlighted GFP localization comparable to the direct fluorescence from GFP, demonstrating that directly incubating antibodies with protoplasts can serve as an alternative IF technique.

**Figure 3.**
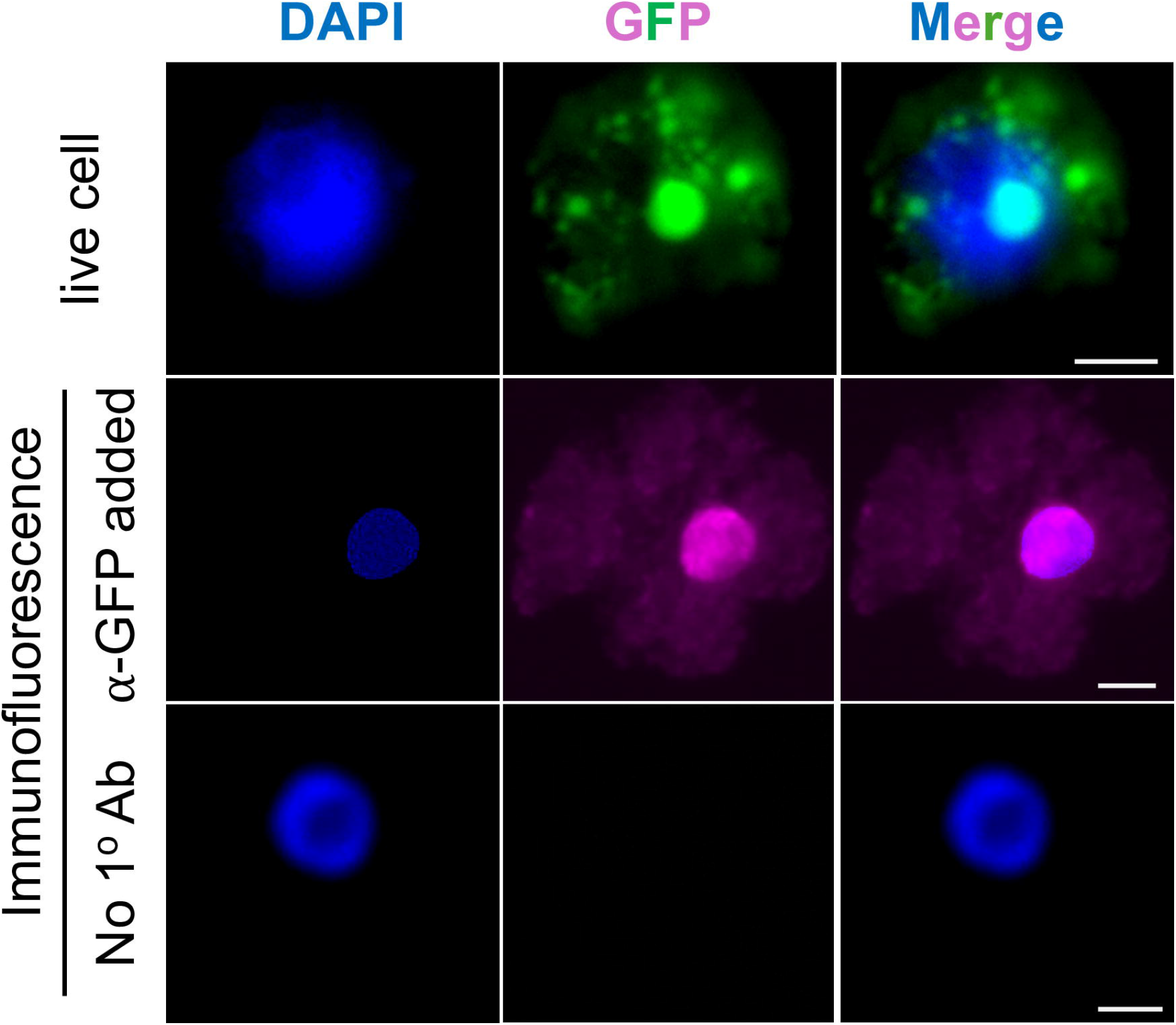
An alternative immunofluorescence protocol. Protoplasts from *Nicotiana benthamiana* line 16C were used for testing. GFP5-ER in protoplasts was detected by ⍰-GFP polyclonal primary antibody and Alexa Fluor594-labeled anti-Rabbit secondary antibody. anti-Rabbit secondary antibody was added to all IF samples. Scale bar, 5 μm.

Using this system, we aimed to understand the subcellular distribution of Pol II and PSTVd RNA. We used the well-known 8WG16 monoclonal antibody to detect the localization of the largest subunit (RPB1) of Pol II. As shown in Figure 4A, there is no fluorescence signal from samples in which we did not add the primary antibody, demonstrating the specificity in detecting target protein. When both antibodies and probes were applied, we observed significant co-localization of PSTVd and Pol II in the nucleoplasm (Figure 4A).

**Figure 4.**
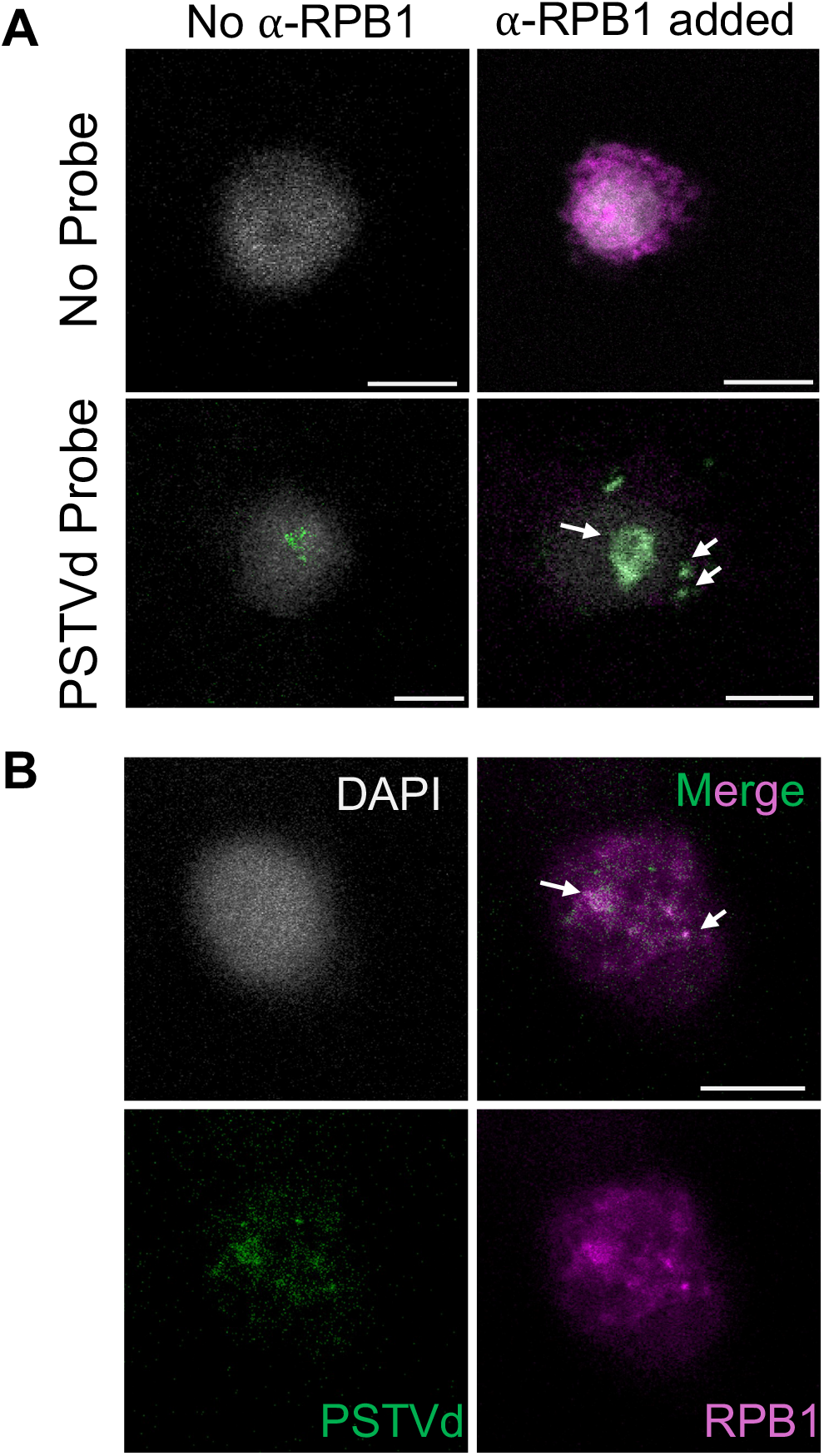
Combining immunofluorescence (IF) and RNA fluorescence in situ hybridization (FISH) to colocalize PSTVd and RPB1. A) RPB1 was detected by ⍰-RPB1 polyclonal antibody and Alexa Fluor594-labeled anti-Rabbit secondary antibody (magenta color). Secondary antibody was applied to all treatments. PSTVd was detected by Alexa Fluor488-labeled specific riboprobe (green). B) Separate color channels of a representative sample showing the colocalization of PSTVd and Pol II (RPB1). White arrow heads point to some colocalized loci. Scale bar, 5 μm.

In another representative image (Figure 4B), we showed that the RPB1 and PSTVd colocalized loci are spread in the nucleoplasm. This pattern demonstrates that Pol II transcribes the template of PSTVd sense-strand RNA in discrete loci mainly in the nucleoplasm. The JACoP analysis showed that Pearson’s co-efficiency values range from 0.867 to 0.997 (Figure S1), indicating a positive co-localization.

### The overall strategy combining IF and FISH to simultaneously observe RNA and cognate binding protein

The success in observing the colocalization of U3 snoRNA/fibrillarin and PSTVd/RPB2 indicates that combined IF-FISH protocol is efficient for observing proteins and RNA simultaneously in the cells (Figure 5). This strategy involves generating protoplasts from leaves, followed by IF process. The key is to fix the cells with antibody before the subsequent RNA FISH steps, which include permeabilization, blocking, hybridization, and finally, observation. The entire procedure can be completed in four days.

**Figure 5.**
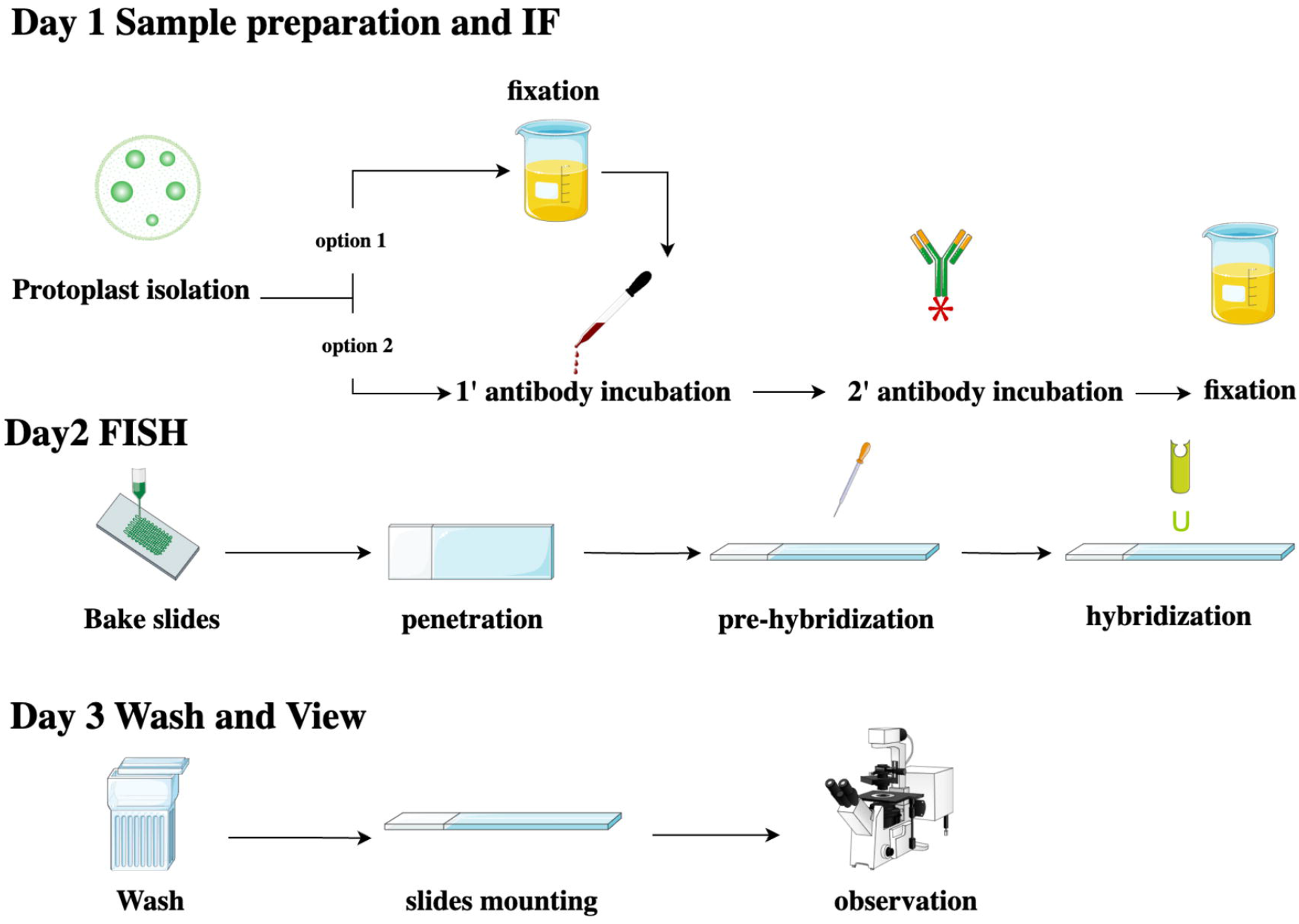
Schematic workflow of combined IF and FISH.

## Discussion

By combining IF and RNA FISH, we successfully achieved the simultaneous visualization of RNAs and their cognate proteins in the nucleus of plant cells. This approach eliminates the need of extensive cloning, as required in the MS2–MCP system (Syed & Lim, 2024), or the generation of transgenic plants when a specific primary antibody is available. Notably, our protocol is effective for detecting both endogenous and foreign RNAs in the nucleus, which has previously proven challenging based on both literature and our experience. Additionally, this method allows the direct visualization of proteins and RNAs in their native state without ectopic expression, making it a robust and straightforward approach for plant cell biology studies.

Compared with smFISH (Duncan et al., 2023), our probe design strategy offers an advantage in detecting relatively short RNAs that are insufficient in length for designing multiple oligos. Nevertheless, our method could be adapted in combination with smFISH to detect endogenous mRNAs. Furthermore, multiple proteins and/or RNAs can be observed within the same cells by utilizing various AlexaFluor dyes, allowing different targets to be marked with distinct colors. This flexibility enables researchers to adapt the method to address a range of biological questions.

Notably, we present two distinct IF protocols for different proteins. The conventional IF protocol is suitable for detecting fibrillarin in our practice. In contrast, the live-cell-antibody-incubation strategy is more suitable for DNA ligase 1 (Wang, Ma, Hao, Liu, & Wang, 2024) and RPB1. It is expected that the two available protocols will increase the chance for successful detection of a wide-spectrum of proteins.

However, we’d like to note that both antibodies and probes penetrate into protoplasts at a relatively low frequency (∼20-30% of cells). Therefore, there are limited cells with both signals. The reasons behind such low frequency are not immediately clear. Nevertheless, it is still possible to find dozens of samples with colocalization signals. Future trials may be needed to further improve the efficiency.

Recent studies have increasingly highlighted the importance of dynamic RNA organization and their associated factors within the cellular environment, particularly in the context of phase separation across diverse biological processes (Miao et al., 2024). However, most knowledge has been derived either from methods lacking cellular context (e.g., RIP-Seq, iCLIP, etc) (Lewinski et al., 2024) or indirectly by observing either protein (Citovsky et al., 2006; Cox et al., 2024; Paciorek, Sauer, Balla, Wisniewska, & Friml, 2006; Sharma, Klosgen, & Bennewitz, 2023) or RNA subcellular localization alone (Duncan, Olsson, Hartley, Dean, & Rosa, 2017; J. Ma et al., 2022; Zollner et al., 2021). Our method provides a means to directly investigate the dynamic interactions between RNAs and proteins in cells, offering a valuable tool for future research. One recent success is demonstrating the co-localization of DNA ligase1 and PSTVd in a biomolecular condensate (Wang et al., 2024).

Although it is well-known that PSTVd replication depends on Pol II and TFIIIA-7ZF activity (Wang et al., 2016), the exact nuclear distribution pattern involved in PSTVd RNA-templated transcription remain unclear. By using our method, we now understand that Pol II transcribes sense-strand of PSTVd in multiple discrete loci in the nucleoplasm. Pol II transcription loci do not all overlap with the major PSTVd biomolecular condensate, and this biomolecular condensate contains PSTVd and DNA ligase1 (Wang et al., 2024), the latter of which is a critical enzyme for generating progeny (Nohales, Flores, & Daros, 2012). Therefore, the observation support the model that there is an active enrichment process of sense-strand PSTVd RNA for enzymatic processing (Qi & Ding, 2003).

## Materials and methods

### Plant growth and protoplast preparation

*Nicotiana benthamiana* (wild type and line 16C) plants were grown in a growth shelf at 25 °C with a 16-hour light and 8-hour dark cycle. PSTVd-infected plants were verified by RNA gel blots as described previously (Takeda, Zirbel, Leontis, Wang, & Ding, 2018). Protoplasts were isolated from the 4^th^ to 6^th^ leaves of the plants by following a published protocol with slight modifications (Jiang, Ma, Liu, & Wang, 2019). Briefly, leaves were digested with 3% cellulose (Onozuka Yakult Pharmaceutical IND., Tokyo, Japan) and 0.8% macerase (MilliporeSigma, Burlington, MA, USA) in protoplast digestion buffer (0.4 M mannitol, 20 mM KCl, 20 mM MES, 10 mM CaCl_2_, 0.1% BSA, 5 mM β-mercaptoethanol, pH 5.7). The lower epidermis layer of the leaves was removed by tapes before digestion. After 1 hr of digestion, the protoplasts were pelleted by centrifugation at 150× *g* for 1 min. The pelleted protoplasts were then washed sequentially with W5 buffer (5 mM MES, 154 mM NaCl, 125 mM CaCl_2_, 5 mM KCl, pH 5.7) at room temperature. Finally, the protoplasts were incubated in the IF buffer (0.5 M mannitol, 20 mM MES, pH 5.7, 20 mM KCl, 1% BSA, 0.1% Tween-20).

### Traditional immunofluorescence (IF)

After extensive trials, we found the following procedures can lead to most clear and repeatable results. Protoplasts were first fixed with 3.7% formaldehyde and 0.2% picric acid in KE buffer (50 mM KH_2_PO_4_, 5 mM EGTA, pH 5.0) at room temperature for 12 min. After fixation, cells were quenched in 1X KE buffer with 300mM glycine for 30 min and washed in 1X KE buffer twice. The fixed cells were then permeabilized in 1X PBS buffer containing 2% BSA, 0.1% cellulase, 0.05% macerozyme, 1% Triton X-100 and 0.5% Nonidet P-40 at room temperature for 30 min, followed by methyl alcohol incubation at minus 20°C for 30 min. Cell blocking was performed using 1X PBS containing 0.2% Tween20, 1% BSA, and 10% DMSO for 1 hr in the room temperature. Cells were incubated with 1:250 diluted fibrillarin primary antibody (Santa Cruz Biotechnology, Dallas, Texas) at 4°C overnight. After three washes, cells were then incubated with 1:250 diluted AlexaFluor594-conjugated anti-mouse secondary antibody (ThermoFisherSci) at 37°C for 1 hr. After three washes, cells were fixed for the second time with 3.7% formaldehyde and 0.2% picric acid in KE buffer at room temperature for 30 min.

### Live-cell immunofluorescence

The live cell IF method was adapted from a study on *in vivo* antibody uptake (Watkins & Burke, 2024). Freshly prepared protoplasts were incubated overnight at 4°C with primary antibodies diluted 1:500 in IF buffer. After primary antibody incubation, protoplasts were washed three times with IF buffer with a 10 min interval. Secondary antibodies were added in IF buffer with a 1:50 dilution and incubated at room temperature for 3 hr. The protoplasts were washed three times with IF buffer (10 min each) to remove the secondary antibody. Sample fixation was done by using 3.7% formaldehyde and 0.2% picric acid in KE at room temperature for 1 hr. Cross-linking was stopped by using 1X KE buffer containing 0.3M Glycine, followed by two times washing with 1X KE buffer for 1h each. Primary antibodies are mouse monoclonal 8WG16 (ThermoFisherSci, Waltham, MA) for Pol II RPB1 and anti-GFP polyclonal antibody raised in rabbit (Genscript, Piscataway, NJ). AlexaFluor594-conjugated anti-mouse or anti-rabbit antibodies (both from ThermoFisherSci) was used as secondary antibody.

### RNA fluorescence in situ hybridization (FISH)

Following IF treatment and fixation, the protoplasts were stored in 70% ethanol at 4°C overnight before FISH steps. The samples were baked onto Superfrost® microscope slides (ThermoFisherSci) at -20°C for 30 minutes, followed by baking at 70°C for 2 hours. Next, the samples were washed with 1X PBS containing 2% bovine serum albumin (BSA), 0.1% cellulase, 0.05% macerozyme, 0.3% Triton X-100, and 0.5% Nonidet P-40 at room temperature for 30 min. Then, samples were washed three times with 100 mM Tris (pH 6.8) containing 1% Nonidet P-40 and 0.5% Triton X-100 at 37°C for 10 minutes each. Pre-chilled methanol with 2% hydrogen peroxide was added to the samples, which were incubated at -20°C for 30 min. The slides were subjected to three rounds of liquid nitrogen treatment, each lasting 5 seconds. The samples were then treated with 0.5% (v/v) acetic anhydride in 0.1 M triethanolamine (TEA) buffer (pH 8.0) at room temperature for 20 min.

Samples were then blocked with PerfectHyb™ Plus hybridization buffer (ThermoFisherSci) at 55°C for 30 min. Probes (corresponding to the 98-188 and 270-13 nucleotide positions in the PSTVd genome but in the complementary orientation) were synthesized using T3 Maxiscript kit (ThermoFisherSci) mixed with AlexaFluor488-labeled UTP (ThermoFisherSci) (J. Ma et al., 2022). Probes (2.5-5 µg/ml) were denatured by heating at 100°C for 1 minutes, followed by cooling on ice. The denatured probes, along with 5 nM ribonucleoside vanadyl complex (Sigma-Aldrich, St. Louis, MO, USA), were added to PerfectHyb™ Plus Hybridization Buffer and applied to samples on slides, followed by incubation at 55°C overnight.

The samples were then rinsed with 2X SSC, washed with 2X SSC containing 50% formamide at 37°C for 30 min, and subsequently washed with 2X SSC and 1X SSC at room temperature for 30 minutes each. 4’,6-diamidino-2-phenylindole (DAPI) was added directly to the samples stain (1 µg/ml) for 5 minutes and then rinsed with distilled water. Anti-fade reagent (ThermoFisherSci) was applied to each sample, and the slides were sealed with nail polish.

### Microscopy imaging

Fluorescence signals were obtained by using a Discover Echo Revolve microscope with DAPI, FITC, and TRITC cubes and Leica Stellaris 5 confocal microscope.

## Acknowledgments

This work is supported by US National Science Foundation (MCB-1906060, MCB-2350392, and IOS-2410009 to YW). Yunhan Wang is supported by CALS Deans’ scholarship from the University of Florida. Symbols in Figure 5 are from bioicons.com. We apologize to colleagues whose work was not cited due to the page limits.

## Notes

### Competing Interest Statement

The authors have declared no competing interest.

### Summary of Updates

We have made extensive updates to the main text and Figures.

