## Supplementary figures and images for "Simultaneous observation of RNAs and their binding proteins in plant cells"

### Figure S1

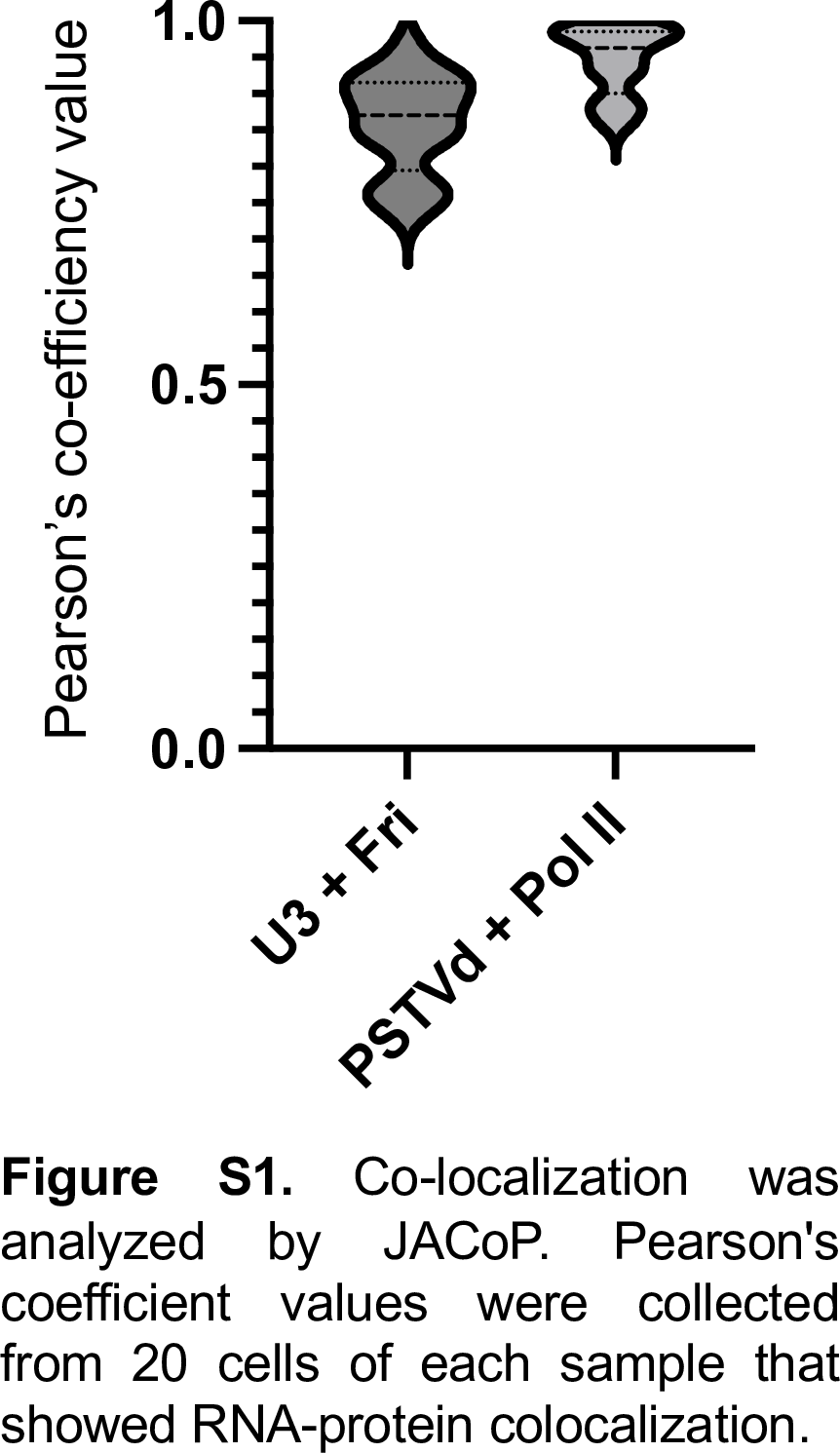
